# Single-cell transcriptomics identifies an effectorness gradient shaping the response of CD4+ T cells to cytokines

**DOI:** 10.1101/753731

**Authors:** Eddie Cano-Gamez, Blagoje Soskic, Theodoros I. Roumeliotis, Ernest So, Deborah J. Smyth, Marta Baldrighi, David Willé, Nikolina Nakic, Jorge Esparza-Gordillo, Christopher G. C. Larminie, Paola G. Bronson, David F. Tough, Wendy C. Rowan, Jyoti S. Choudhary, Gosia Trynka

**Author notes:** These authors contributed equally to this work (Eddie Cano-Gamez, Blagoje Soskic). Correspondence to: Gosia Trynka and Blagoje Soskic.

## Abstract

Naïve CD4+ T cells coordinate the immune response by acquiring an effector phenotype in response to cytokines. However, the cytokine responses in memory T cells remain largely understudied. We used quantitative proteomics, bulk RNA-seq and single-cell RNA-seq of over 40,000 human naïve and memory CD4+ T cells to generate a detailed map of cytokine-regulated gene expression programs. We demonstrated that cytokine response differs substantially between naïve and memory T cells and showed that memory cells are unable to differentiate into the Th2 phenotype. Moreover, memory T cells acquire a Th17-like phenotype in response to iTreg polarization. At the single-cell level, we demonstrated that T cells form a continuum which progresses from naïve to effector memory T cells. This continuum is accompanied by a gradual increase in the expression levels of chemokines and cytokines and thus represents an effectorness gradient. Finally, we found that T cell cytokine responses are determined by where the cells lie in the effectorness gradient and identified genes whose expression is controlled by cytokines in an effectorness-dependent manner. Our results shed light on the heterogeneity of T cells and their responses to cytokines, provide insight into immune disease inflammation and could inform drug development.

## Introduction

A healthy immune system is characterized by efficient communication between cells, which facilitates a quick response to a wide variety of pathogens. This communication is mediated by cytokines. Upon binding to their receptors, cytokines trigger a signaling cascade which culminates with the induction of gene expression programs^1, 2^. This promotes the differentiation of target cells into effector cell types. This process is particularly relevant for CD4+ T cells, which coordinate the downstream response of various immune cells (e.g. CD8+ T cells, macrophages and B cells)^3^. Triggering of the T cell receptor (TCR) and co-stimulatory molecules activates naïve CD4+ T cells, which are then directed by cytokines to polarize into various T helper (Th) phenotypes. These include Th1, Th2 and Th17, which secrete IFN-γ, IL-4 and IL-17, respectively^4–7^. Moreover, in response to transforming growth factor beta (TGF-β), naïve CD4+ T cells acquire regulatory potential (induced regulatory T cells, iTreg) and suppress effector T cell responses^8^.

Previous *in vitro* studies investigated how cytokines modulate T cell function^9–18^, increasing our understanding of cytokine-induced polarization. Nonetheless, most studies have focused exclusively on naïve CD4+ T cells, altogether excluding memory cells. This is in part due to the premise that, once T cells undergo stimulation and respond to a cytokine, the phenotype acquired by CD4+ T cells remains mostly stable. Recent studies have challenged this idea, providing evidence that cytokines can reprogram the phenotypes of polarized T cells^2, 19, 20^. For example, IL-6 can convert Treg cells to a pathogenic Th17-like phenotype under arthritic conditions^21^. Furthermore, Th17 cells upregulate *TBX21* and IFN-γ in response to Th1-polarizing cytokines^22^, and infection-induced Th17 cells from the gut can secrete a variety of inflammatory cytokines, e.g. the Th1 cytokine IFNγ^23^. These observations highlight the remarkable plasticity of CD4+ T cells and suggest that memory cells retain the ability to respond to cytokines. However, understanding the effects of cytokines on memory T cells is challenging because circulating memory T cells are heterogeneous, comprised of multiple subpopulations such as central and effector memory cells^24–26^.

Cytokines also play a central role in autoimmunity and are often tractable, and successful therapeutic targets. Twenty-five years ago, injectable IFN-β was approved as the first disease modifying therapy (DMT) for multiple sclerosis^27^, yet the therapeutic mechanism is still unknown. Another DMT for multiple sclerosis is an immune modulator which shifts the cytokine profile of pro-inflammatory Th1 cells to anti-inflammatory Th2 cells^28^. These observations are not yet fully understood and illustrate how increasing our understanding of cytokine responses is crucial for improved drug development.

Finally, genetic studies have implicated CD4+ T cells, and particularly memory T cells, in the biology of many common complex immune diseases^29–31^, suggesting that it is especially relevant to understand cytokine responses in memory T cells. Therefore, limiting the study of cytokines to the naïve T cell compartment could bias our understanding of the processes underlying pathologic inflammation.

In this study, we characterized the response of naïve and memory CD4 T cells to five different cytokine conditions influencing inflammation and immune diseases. To account for the dynamic nature of cytokine responses, we profiled cells at two different time points following stimulation. We also examined cells in the resting state, resulting in a total of 12 distinct cell states. We used these profiles to generate a detailed map of gene and protein expression changes induced by cytokines. This map leverages information from quantitative proteomics, RNA-sequencing and single-cell RNA-sequencing (scRNA-seq) of over 40,000 single T cells, thus providing a comprehensive resource with exceptional resolution. We found that naïve T cells responded differently to cytokines than memory T cells. At the single-cell level, we recapitulated previously described CD4+ T cell subpopulations and found that T cells did not form discrete groups of cells but instead formed a continuum characterized by a gradual increase in the expression level of chemokines, granzymes and other effector molecules. Importantly, this gradient was present in the resting state, persisted after stimulation and determined how cells respond to cytokines by modulating the magnitude of cytokine-induced gene expression changes.

## Results

### Study design

To investigate the effects of cytokines on the two main subsets of human CD4+ T cells, we purified CD4+ CD25− CD45RA+ CD45RO-naïve T (T_N_) cells and CD4+ CD25− CD45RA− CD45RO+ memory T (T_M_) cells (**Supplementary Figure 1A** and **Methods**). We then stimulated the cells with anti-CD3/anti-CD28 coated beads in the presence of different cytokine cocktails (**Figure 1A**, **Figure 1B** and **Supplementary Table 1**). We selected cytokine cocktails to polarize T_N_ and T_M_ cells towards four major T helper phenotypes (Th1, Th2, Th17 and iTreg). In addition, we included IFN-β due to its role as a therapy in multiple sclerosis^32, 33^. In order to distinguish T cell responses to TCR/CD28-activation from responses induced specifically by cytokines, we stimulated cells with anti-CD3/anti-CD28 beads in the absence of any cytokines (Th0). Finally, we also cultured cells without neither stimulation nor cytokines (resting cells). We profiled gene expression (RNA-seq) for early transcriptional responses (16 hours after stimulation, before cell proliferation) and late transcriptional responses (5 days after stimulation, after cell proliferation), when cells are thought to acquire an effector phenotype. To comprehensively characterise cellular states at the late time point, we also profiled the whole proteome using isobaric labelling with two-dimensional liquid chromatography-tandem mass spectrometry (LC-MS/MS), as well as the transcriptome at the single cell level (scRNA-seq) (**Methods**).

**Figure 1.**
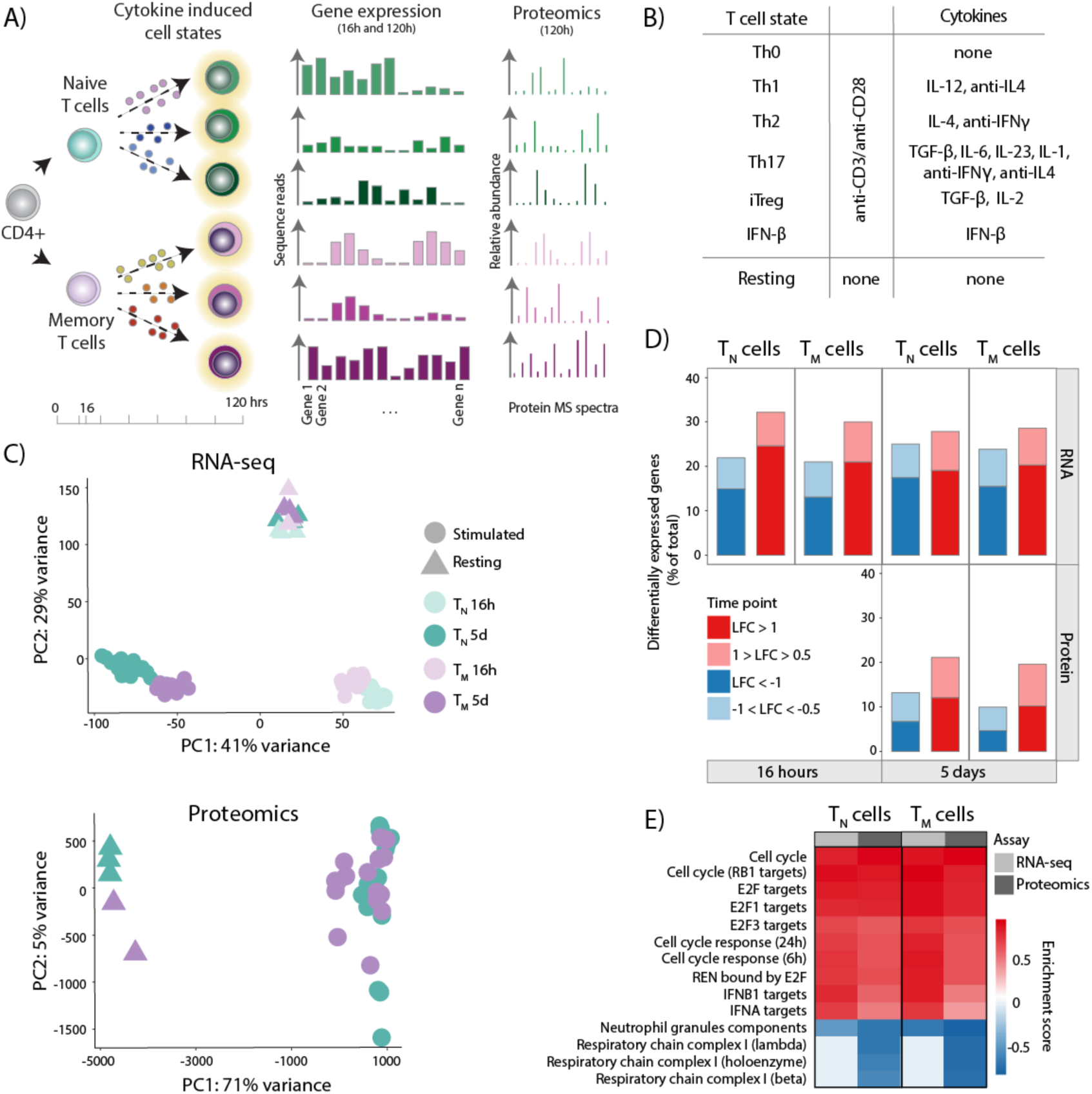
TCR/CD28-activation induces cell type specific gene expression programs in CD4^+^ T cells. **A)** Overview of the experimental design. **B)** List of cytokine conditions. **C)** PCA plots from the whole transcriptome (upper panel) and proteome (lower panel) of T_N_ and T_M_ cells. Different colors correspond to cell types and different shades to stimulation time points. **D)** Gene expression changes at the RNA and protein levels by comparing TCR/CD28-activated (Th0) cells to resting cells. Up-regulated genes are in red and down-regulated genes are in blue. Different shades indicate different fold-change thresholds. **E)** A selection of significantly enriched pathways (with enrichment scores > 0.7) from genes and proteins differentially expressed after five days of activation using the 1D enrichment method.

### TCR/CD28-activation induces well-defined gene expression programs in naïve and memory T cells

To understand T_N_ and T_M_ cell responses to T cell activation (TCR/CD28-activation), we compared the transcriptomes of activated and resting cells. We observed that the main source of variation across the full transcriptome and proteome was T cell activation, with resting cells clustering separately from activated cells (**Figure 1C**). Activated cells also clustered by duration of stimulation (16 hours and five days) and cell type (T_N_ and T_M_), suggesting that the response to T cell stimulation is dynamic and cell type specific (**Figure 1C**). We then tested for differential RNA and protein expression between resting and activated (Th0-stimulated) T_N_ and T_M_ cells. We identified a large number of changes which develop in response to stimulation (**Figure 1D, Supplementary Tables 2** and **Supplementary Table 3**). At the RNA level, 8,333 and 7,181 genes (corresponding to approximately 40% of the transcriptome) were differentially expressed after 16 hours of activation in T_N_ and T_M_ cells, respectively. This number was comparable after five days (7,705 and 7,544 in T_N_ and T_M_ cells). At the protein level, we identified 4,009 and 3,443 differentially expressed proteins (approximately 35% of the proteome data) after five days of activation in T_N_ and T_M_ cells, respectively. These genes formed a well-defined expression program characterized by upregulation of the cell cycle and targets of the E2F family of transcription factors, as well as the type I IFN response (**Figure 1E** and **Supplementary Table 4**). Conversely, T_N_ and T_M_ cells downregulated components of the respiratory chain complex in response to activation (**Figure 1E**). This is in line with previous observations suggesting that T cell activation induces proliferation and profound metabolic changes to support effector responses^34^. Importantly, these conclusions were consistent between RNA and protein.

### Cytokines induce cell type specific gene expression programs in naïve and memory CD4+ T cells

We next investigated how cytokines modulate gene expression in T_N_ and T_M_ cells. We first performed PCA on the full proteome and transcriptome, treating time points and cell types independently. While there were few cytokine effects at 16 hours (**Supplementary Figure 1B**), we observed clear clustering by cytokine condition at five days (**Figure 2A**) that were consistent between transcriptome and proteome. To disentangle cytokine effects from those of T cell activation (TCR/CD28-activation), we compared stimulated cells exposed to cytokines to Th0-stimulated cells (**Supplementary Table 2** and **Supplementary Table 3**). Most cytokine induced changes were only apparent after five days of stimulation (**Figure 2B**), with the exception of IFN-β. For example, Th17-stimulation induced only 42 differentially expressed genes after 16h in naïve cells at the RNA level, compared to 1,818 differential genes induced after 5 days. This pattern was similar for other cytokine conditions. In contrast, IFN-β induced a large number of early transcriptional changes (357 genes at 16h and 329 after 5 days in T_N_ cells), reflecting its’ role in the fast response to viruses. These results suggest that early changes in gene expression are dominated by the effects of T cell activation alone, while the expression programs characteristic of differentiated Th cells are apparent at the later stages of stimulation. This implies that cytokine polarization occurs not in parallel but after the initiation of T cell activation.

**Figure 2.**
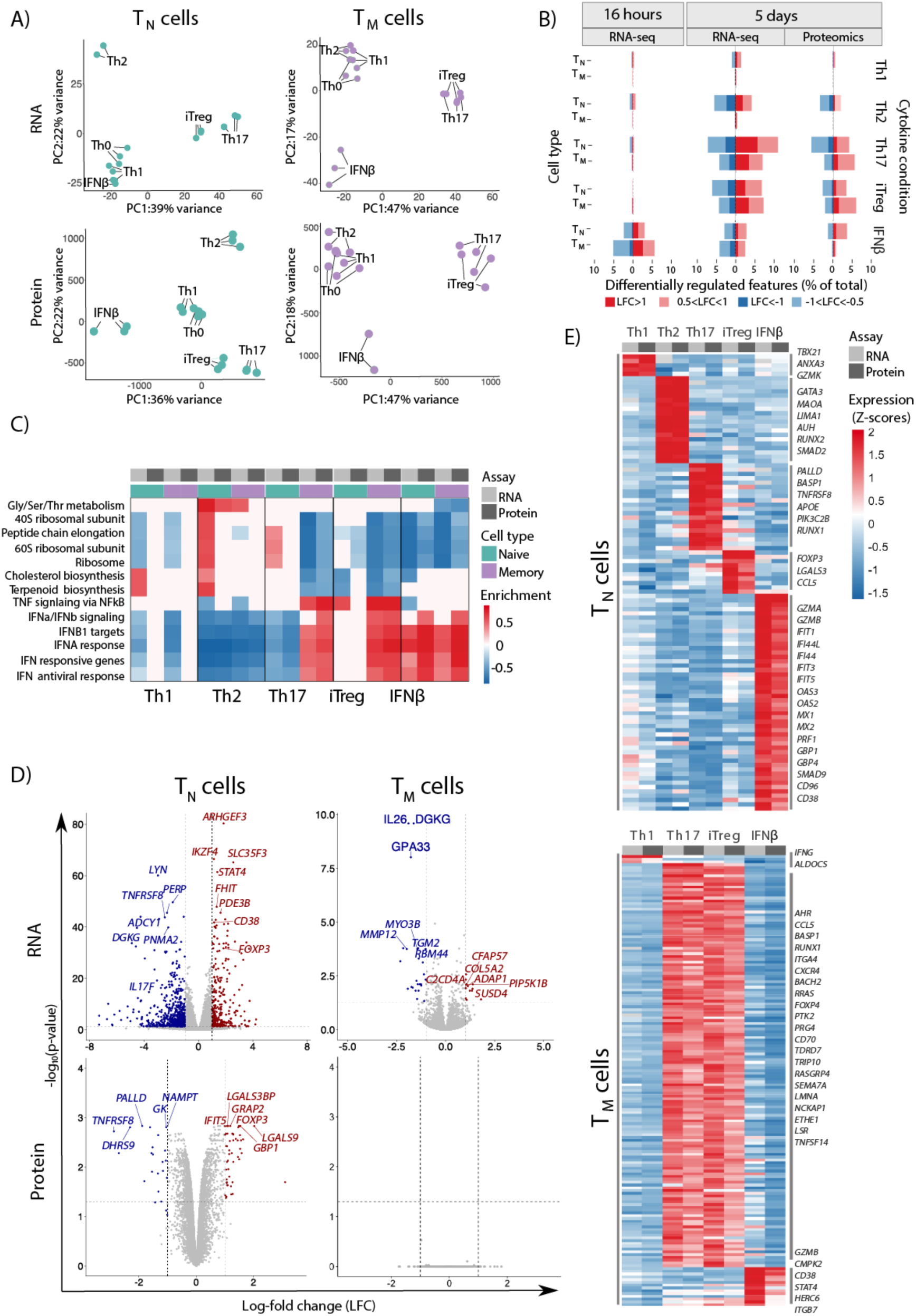
Cytokines induce cell type specific gene expression programs in CD4^+^ T cells. **A)** PCA plot from the full transcriptome and proteome of T_N_ and T_M_ cells following five days of cytokine stimulations. Only stimulated cells were included in this analysis. **B)** Gene expression changes at the RNA and protein levels from pairwise comparisons between cytokine-stimulated cells and Th0-stimulated cells. Up-regulated genes are in red and down-regulated genes are in blue. Different shades indicate different fold-change thresholds. **C)** A selection of significantly enriched pathways (with enrichment scores > 0.7) from differentially expressed genes and proteins using the 1D enrichment method. **D)** Volcano plots highlighting significant differences in gene and protein expression between Th17 and iTreg-stimulated T_N_ and T_M_ cells. Red indicates expression upregulation in iTreg with respect to Th17-stimulation, blue indicates expression upregulation in Th17 with respect to iTreg-stimulation. Labels were added to *IL17*, *FOXP3* and the top 20 most differentially expressed genes. **E)** Cell state specific gene signatures defined using jointly RNA and protein expression. Colours encode normalized (Z-scored) gene and protein expression levels. Example genes for each signature are labeled.

Since cytokine induced effects mainly manifested five days after stimulation, we subsequently focused on the late time point to further elucidate changes in gene and protein expression driven by different cytokines. We then compared these effects between T_N_ and T_M_ cells. In general, the number of cytokine-induced changes in RNA and protein expression was comparable between T_N_ and T_M_ cells (**Figure 2B**). However, Th2-stimulation clearly triggered different responses between the two cell types, resulting in differential expression of 944 genes in T_N_ cells compared to 49 in T_M_ cells. We observed the same trend at the protein level, where 290 proteins were differentially expressed in T_N_ cells but no differences were detected in T_M_ cells in response to Th2-stimulation (**Figure 2B**). This was true despite T_M_ cells expressing comparable levels of the IL-4 receptor than T_N_ cells (**Supplementary Figure 2A**). This suggested that T_M_ cells cannot be polarized towards the Th2 phenotype.

We next sought to translate these observations to cellular functions and pathways. We observed that the genes and proteins differentially expressed upon cytokine stimulation formed well-defined expression programs and were enriched in relevant pathways (**Figure 2C** and **Supplementary Table 4**). As expected, stimulation with IFN-β induced upregulation of the type I IFN response in both T_N_ and T_M_ cells, while Th2-polarization of T_N_ cells suppressed this pathway, likely reflecting that Th2-polarization involves IFN-γ blockade. Importantly, these effects were concordant between RNA and protein. Furthermore, we found that Th1-stimulation of T_N_ cells induced metabolic changes such as increased cholesterol and terpenoid synthesis, while Th2-stimulation increased the expression of genes involved in amino acid metabolism (**Figure 2C**). Interestingly, some pathways showed opposite effects between T_N_ and T_M_ cells upon cytokine stimulation. For example, while Th17-stimulation of T_N_ cells induced downregulation of the type I IFN response, Th17-stimulation of T_M_ cells increased the activity of this same pathway. We observed a similar pattern upon iTreg-stimulation, with the type I IFN response being upregulated in T_M_ but not in T_N_ cells (**Figure 2C**). These observations suggest that Th17 and iTreg-stimulation conditions induce different cell states in T_N_ than in T_M_ cells.

Th17 and iTreg cells have been extensively linked to autoimmune inflammation and immune suppression. Polarization to both of these cell states requires the presence of TGF-β and there is evidence of interconversion between the two cell phenotypes^21^, suggesting that their functions may be interrelated. Consistent with these observations, we found that Th17 and iTreg-stimulated T_N_ and T_M_ cells were more similar to each other than to any other cell state and formed a single cluster on the PCA plot (**Figure 2A**). This similarity was captured by both proteome and transcriptome. This was in sharp contrast with T_N_ cells, where the two cytokine-induced cell states formed separate groups. Importantly, both cell types expressed comparable levels of the TGF-β and IL2 receptors (**Supplementary Figure 2B** and **Supplementary Figure 2C**). To further test whether Th17 and iTreg-stimulation induced the same phenotype in T_M_ cells, we compared the expression of genes between the two cell states (**Figure 2D, Supplementary Table 2** and **Supplementary Table 3**). Only 42 genes and no proteins were differentially expressed between the two cytokine conditions in T_M_ cells at the selected thresholds (LFC > 1 at 0.05 FDR for RNA-seq and LFC > 0.5 at 0.1 FDR for proteomics). In contrast, in T_N_ cells 733 genes and 455 proteins were differentially expressed between iTreg and Th17-stimulated cells (**Figure 2D**). In particular, iTreg-stimulated T_N_ cells expressed higher levels of *FOXP3*, *IKZF4* and *LGALS3*, while Th17-stimulated T_N_ cells expressed higher levels of *IL17F*, *TNFRSF8* and *PALLD*. Therefore, while T_N_ cells acquire different phenotypes upon Th17 and iTreg-polarization, both cytokine conditions polarize T_M_ cells towards the same cell state.

### Cell state specific gene signatures from proteome and transcriptome

Our results suggested that these cytokines act in a cell type specific manner to induce five well-defined cell states in T_N_ cells (Th1, Th2, Th17, iTreg and IFN-β) and three well-defined cell states in T_M_ cells (Th1, Th17/iTreg and IFN-β, while lack Th2 response). We next set out to identify the most specific genes characterizing these cell states. We confirmed that RNA and protein expression showed high correlation in our data, both at the sample and at the gene level (**Supplementary Figure 1C and 1D**). Thus, we applied a multi-omics approach which leveraged both layers of molecular information to derive robust cell state gene signatures (**Methods**). This approach allowed us to identify genes with concordant effects in the two assays, thus increasing our confidence that we captured true cytokine-induced effects. In brief, we identified differentially expressed RNA-protein pairs and asked if any of these pairs were present at a higher level in one cell state compared to the rest. In this way, we derived a measurement of cell state specificity for each gene, where genes with higher specificity than expected by chance were included in a cell state specific *proteogenomic signature* (**Methods**). Because we derived these signatures jointly from transcriptome and proteome, our signatures were sensitive to relative changes in both RNA and protein levels.

In T_N_ cells we identified 105 signature genes corresponding to one of the five different cell states (5 genes for Th1, 20 for Th2, 20 for Th17, 10 for iTreg, and 50 for IFN-β) (**Figure 2E** and **Supplementary Table 5**). The T_N_ IFN-β signature contained well established antiviral genes involved in IFN-regulated functions such as RNAse L induction (*OAS2*, *OAS3*), GTPase activity (*MX1*, *MX2*) and cell lysis (*GZMA*, *GZMB*)^35^. Signatures of other T_N_ cell states also included well known hallmark genes, such as *GATA3* (Th2 signature), *TBX21* (Th1 signature) and *FOXP3* (iTreg signature) (**Figure 2E**). This illustrated that our approach accurately identified known markers of cytokine polarization. Moreover, we observed genes highly specific to Th1 (*ANXA3*), Th2 (*MAOA, LIMA1, MRPS26*), Th17 (*TNFRSF8, RUNX1, PALLD*) and iTreg (*LMCD1, LGALS3, CCL5*) cell states which have not been previously described in the context of cytokine polarization.

We performed the same analysis for T_M_ cells, where we identified 162 signature genes corresponding to one of the three cell states (three for Th1, 145 for Th17/iTreg and 14 for IFN-β genes) (**Figure 2E** and **Supplementary Table 5**). Since Th17 and iTreg-stimulated T_M_ cells overlapped on both the RNA and protein levels, we treated them as one phenotype in this analysis. We observed that the IFN-β signature was substantially different in T_M_ compared to T_N_ cells (only 6 genes overlapped between both signatures). Nonetheless, it contained well known antiviral genes such as *HERC6* and *GZMB*. Several Th17/iTreg T_M_ signature genes were also present in the iTreg and Th17 signatures derived from T_N_ cells (*CCL5, LGALS3, TNFRSF8*) or had been previously linked to one of the two phenotypes in the literature (*BACH2, BATF3, AHR*), suggesting that Th17/iTreg T_M_ cells might have overlapping functions with the Th17 and iTreg states in T_N_ cells. The signature genes identified with our approach provide a valuable resource for future follow-up studies in specific biological contexts or disease settings.

### Single-cell transcriptomics reveals a CD4+ T cell effectorness gradient

Our results showed that the gene expression programs induced in response to cytokines can differ substantially between T_N_ and T_M_ cells. While T_N_ constitute a rather uniform cell population, T_M_ cells are composed of multiple subpopulations including central (T_CM_) and effector (T_EM_) memory cells, as well as effector memory cells re-expressing CD45RA (T_EMRA_). Given this heterogeneity, we speculated that the observed differences in cytokine responses could be explained in two ways: i) T_M_ cells, as a whole, are unresponsive to certain cytokines, or ii) specific subpopulations of T_M_ cells respond to cytokines but the measured bulk gene expression profiles are dominated by a large proportion of unresponsive cells. To address this, we profiled single-cell gene expression in a total of 43,112 T_N_ and T_M_ cells, which included resting cells, as well as cells exposed to Th0, Th2, Th17 and iTreg-stimulation.

First, we isolated T_N_ and T_M_ cells from four healthy individuals and quantified gene expression in the resting state using droplet-based single-cell RNA-sequencing^36^ (**Methods** and **Supplementary Figure 3A**). In total, we profiled 5,269 resting T cells (2,159 T_N_ and 3,110 T_M_ cells respectively), with an average of 1,146 genes detected per cell. We identified 64 highly variable genes, which we used for dimensionality reduction and embedding with the uniform manifold approximation (UMAP)^37^, as well as for unsupervised cell clustering (**Methods**). We identified five distinct groups of cells (**Figure 3A**) which we annotated as T_N_, T_CM_, T_EM_, T_EMRA_ and natural T regulatory (nTreg) cells based on the expression of well established cell type markers (**Figure 3B** and **Supplementary Table 6**). T_EMRA_ cells showed a distinct transcriptional profile characterized by high expression of cytotoxic genes (eg. *PRF1*, *CCL4*, *GZMA*, *GZMH*), consistent with previous observations^25^. Importantly, these cells expressed comparable levels of CD4 and no CD8. The proportions of cells detected in each cell subpopulation were comparable across all biological replicates (**Figure 3A**). We observed that T_N_ cells were mostly a homogeneous group of cells. However, a small percentage of T_EMRA_ cells were originally isolated as T_N_ cells (as they re-express the CD45RA marker) and could only be correctly identified at the single-cell level.

**Figure 3.**
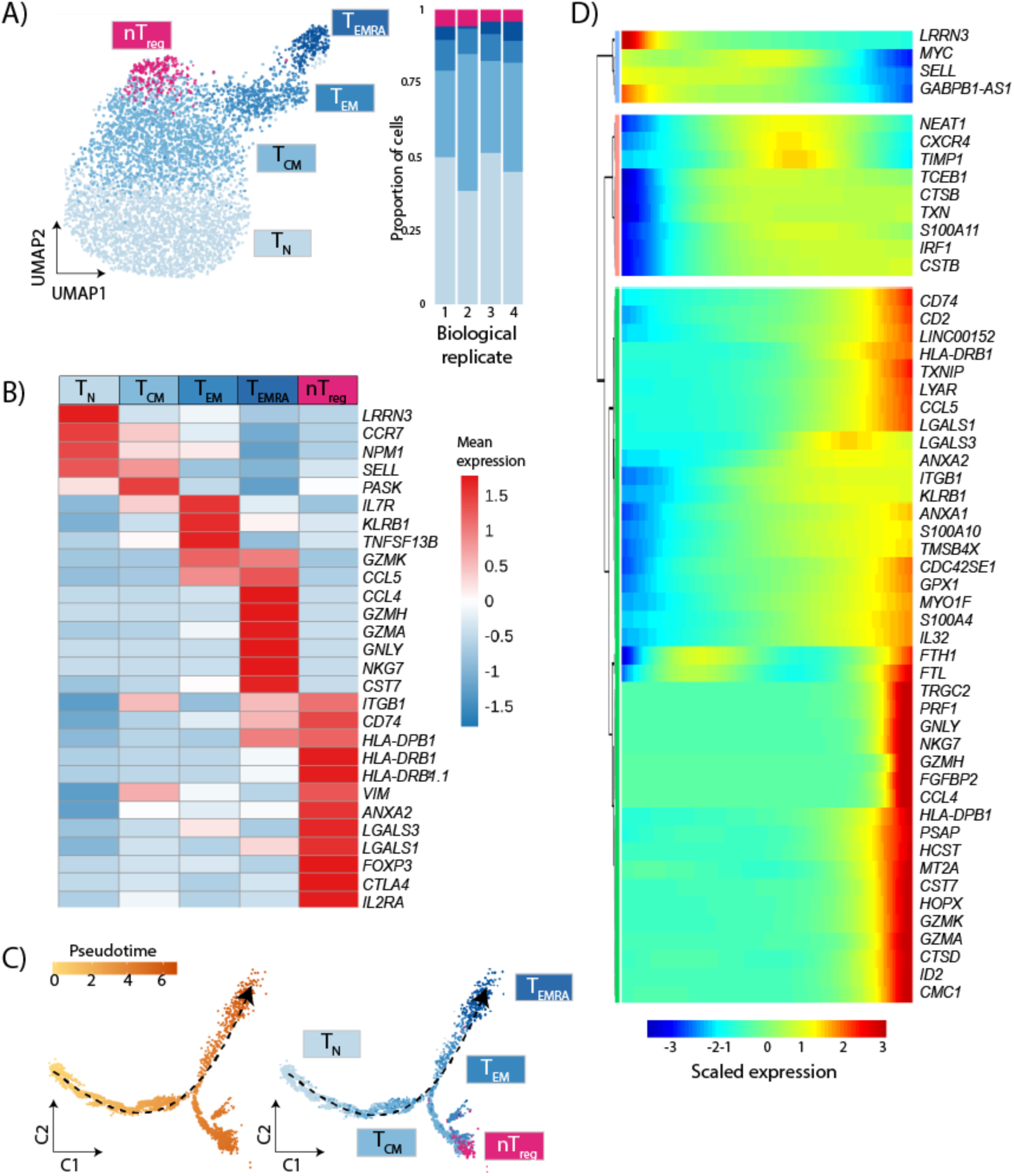
Effectorness gradient in resting CD4+ T cells. **A)** UMAP of single-cell RNA-seq data from resting T cells. Colors represent cells in the five clusters defined using top variable genes and unsupervised clustering. Bar plots represent the proportion of cells assigned to different clusters in each biological replicate. **B)** Gene markers of each cell cluster (Wilcoxon rank sum test) combined with well known markers from the literature. Colors encode the mean expression of each gene in each cluster. **C)** Branched pseudotime trajectory, each cell is colored by its pseudotime value (left panel) or its cluster label (right panel), as determined in panel A. **D)** Heatmap of genes variable along the pseudotime trajectory (from Monocle). The X axis represents cells ordered by pseudotime (from left to right) and different colors correspond to the scaled (Z-scored) expression of each gene in each cell.

In addition, our results suggested that CD4+ T cells do not consist of discrete subpopulations. Instead, different subsets of T cells localized to different areas of the same population within the UMAP space (**Figure 3A**), suggesting that they form one population with multiple interrelated transcriptional states. To investigate the relationships between these states we applied pseudotime analysis^38^ to determine if the cells formed a continuous trajectory. We identified a clear trajectory, starting with T_N_ cells and gradually progressing towards T_M_ cells. The cells at the beginning of this trajectory expressed high levels of naïve markers (e.g. *CD62L, CCR7* and *LRRN3*). In contrast, the end of the trajectory was enriched in cells expressing high levels of cytotoxic molecules (e.g. *GZMA, GZMB* and *PRF1*). This approach confirmed that transcriptionally CD4+ cells formed a natural progression from the least effector (T_N_) to the most effector (T_EMRA_) cell subset (**Figure 3C**), with nTreg cells branching separately from the main trajectory. Furthermore, the expression of cytokines and chemokines, such as *IL32*, *CCL4* and *CCL5*, gradually increased along the pseudotime axis (**Figure 3D** and **Supplementary Table 7**). Therefore, this pseudotime ordering corresponded to the levels of T cell effector functions, which formed a gradient. We refer to this as *effectorness*. Taken together, our results demonstrate that CD4+ T cells are a continuum of cells with varying effectorness, rather than a collection of discrete cell subsets.

### Single-cell transcriptomics separates cells by effectorness and cytokine-induced cell state

Given the observed effectorness gradient we next assessed if this influenced cell responses to T cell activation and cytokine polarization. Therefore, we exposed T_N_ and T_M_ cells to Th0, Th2, Th17 and iTreg-stimulation and profiled single-cell gene expression five days following stimulation. We combined the data obtained from these four conditions into a single data set, which contained single cell transcriptomic profiles of 37,843 cells, of which 18,786 were T_N_ and 19,057 were T_M_ cells, respectively. We used these single-cell profiles to identify 220 highly variable genes, which were used for dimensionality reduction and UMAP embedding. We observed that T_N_ and T_M_ cells formed one single cluster of cells but separated into two different areas of the UMAP space (corresponding to UMAP1, **Figure 4A**), which is in agreement with our observations from resting T cells. We also found that cells exposed to different cytokines localized to different areas of the UMAP space (**Figure 4B**). This was confirmed by a high expression of literature markers associated with the respective cytokines (**Figure 4C**). For example, iTreg-stimulated T_N_ cells localized to an area with high expression of *CTLA4*, while the area associated with Th17-stimulated T_N_ cells showed high *RORA* expression. The area enriched in Th17-stimulated T_M_ cells showed higher levels of *IL17F* (**Figure 4C**). Moreover, cells in these areas also showed higher expression of the corresponding genes identified from our proteogenomic signature analysis (**Supplementary Figure 3B**), confirming a high overlap between our bulk and single-cell observations. Importantly, while the response of T_N_ cells to cytokines was homogeneous, T_M_ cells exposed to cytokines fragmented into multiple groups (**Figure 4B**), suggesting the existence of different gene expression programs specific to T_M_ subpopulations.

**Figure 4.**
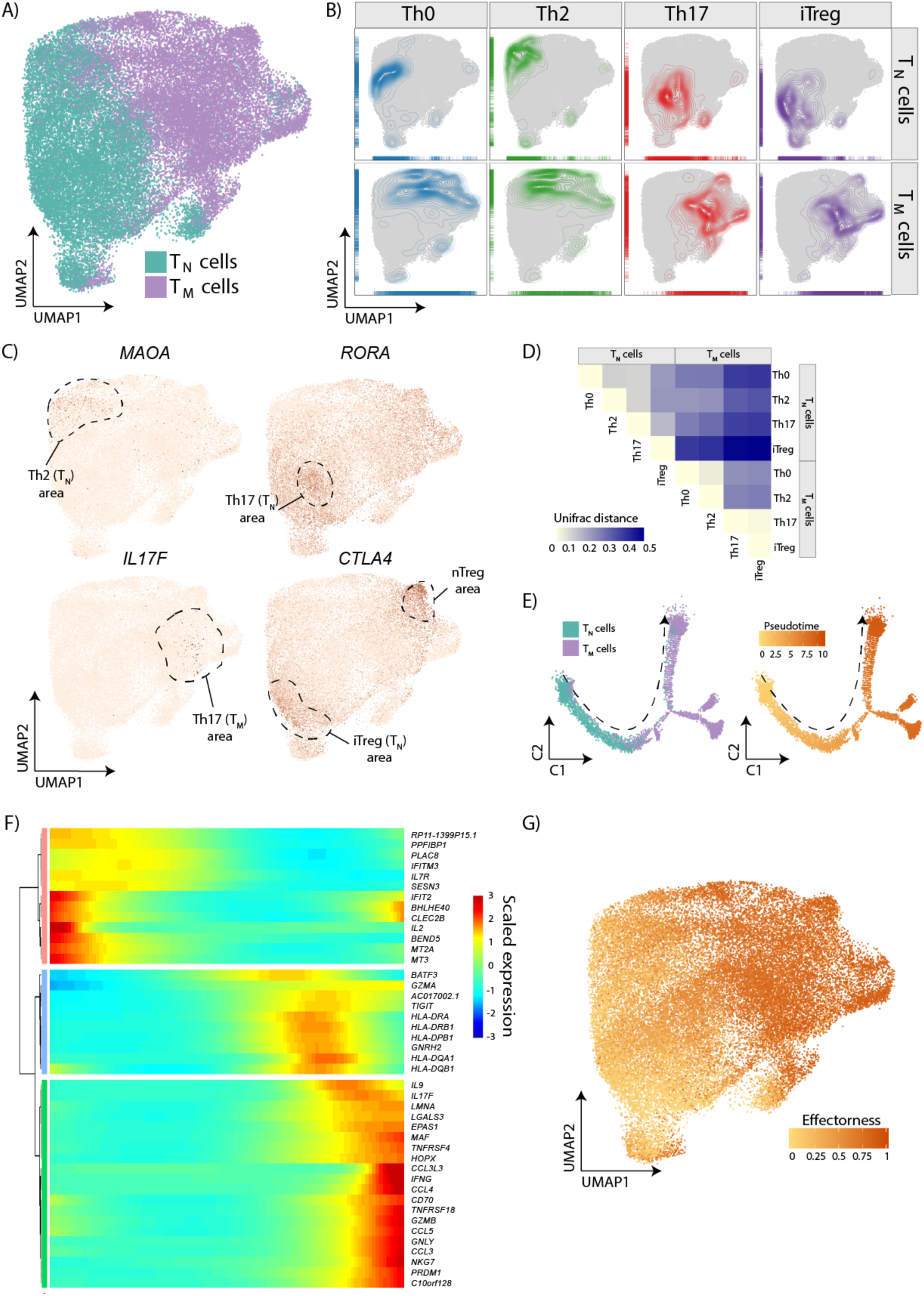
CD4+ T cells separate by effectorness and cytokine-induced cell state. **A)** UMAP embedding of single stimulated T cells into a two-dimensional space. Green corresponds to T_N_ and purple to T_M_ cells. **B)** Density plots highlighting cells based on the cytokines they were exposed to. **C)** Expression of cytokine markers described in the literature. Each dot represents a single cell and colors correspond to the expression level of a gene in each cell. **D)** UniFrac distances between T_N_ and T_M_ cells exposed to different cytokines summarized in a correlation plot. **E)** Th0-stimulated T_N_ and T_M_ cells ordered in a branched pseudotime trajectory. Each cell is colored by cell type (left panel) or pseudotime value (right panel). **F)** Heatmap of the most variable genes along the Th0 pseudotime trajectory (from Monocle). The X axis represents cells ordered by pseudotime (from left to right) and colors correspond to the scaled (Z-scored) expression of each gene in each cell. **G)** An overlay of the cells’ *effectorness* values into the UMAP embedding described in panel A. Cells exposed to each cytokine condition were ordered in separate branched pseudotime trajectories using Monocle, these trajectories were subsequently combined into a single numeric variable called *effectorness*.

Based on the observations from bulk gene expression, we asked whether the absence of response to Th2-stimulation in T_M_ cells was characteristic of the entire population of cells or if a specific subpopulation responded to Th2-stimulation but was masked by a majority of unresponsive cells. Interestingly, Th0 and Th2-stimulated T_M_ cells predominantly localized to the same UMAP areas (**Figure 4B**). We used UniFrac distances^39^, to formally test if Th0 and Th2-stimulated T_M_ cells overlapped (i.e. localized to the same clusters) or formed different groups. The UniFrac method first groups cells in a dendrogram based on their transcriptome. Next, it compares the average position of cells from different samples in the dendrogram and summarizes these differences in a single distance metric^40^. A UniFrac distance of 0 indicates that cells from the two groups have exactly the same composition, while a distance of 1 indicates that the groups form entirely separate clusters. We confirmed that Th0 and Th2-stimulated T_M_ cells nearly perfectly overlapped (UniFrac distance = 0.047) (**Figure 4D**) indicating that no individual subpopulations of T_M_ cells were capable of responding to Th2-stimulation. Instead, the observed lack of response was a uniform characteristic of all T_M_ cells.

Our observations from the bulk data also support that T_M_ cells polarize to the same cell state in response to both Th17 and iTreg-stimulation. We confirmed this at the single cell level, where cells from these two conditions localized to the same UMAP areas. The UniFrac distance between these cell states was 0.015 in T_M_ cells, compared to 0.164 in T_N_ cells (**Figure 4D**). Thus, we concluded that in response to Th17 and iTreg-stimulation T_M_ cells converge on the same cell state. This is not driven by any subpopulation of T_M_ cells and is rather a general characteristic of memory T cell biology. Interestingly this population expressed high levels of *IL17F,* suggesting that iTreg-stimulation in T_M_ cells induces a Th17-like phenotype.

We next assessed how the effectorness gradient affected cell response to stimulation in the presence of cytokines. We observed that the most widely used markers of central and effector memory T cells (i.e. CD62L and CCR7) significantly changed in expression patterns following activation (**Supplementary Figure 3C**), making the annotation of individual T_M_ subpopulations challenging. To overcome this, we applied pseudotime ordering to infer the effectorness of each single cell. In brief, we ordered cells within each cytokine condition into branched pseudotime trajectories. This resulted in four cytokine-specific pseudotime trajectories (Th0, Th2, Th17 and iTreg). First, we analyzed Th0-stimulated cells and confirmed that the trajectory inferred for this cell state showed a similar pattern to that observed in resting cells (**Figure 4E** and **Supplementary Table 7**). We observed a gradual increase in the expression of cytokines and effector molecules which progressed from T_N_ to T_M_ cells. A similar pattern was apparent in other cytokine stimulations, with T_N_ cells localizing to the beginning and T_M_ cells to the end of the respective trajectories (**Supplementary Figure 4** and **Supplementary Table 7**). Furthermore, the end of these trajectories contained cells expressing high levels of genes specific to T_EMRA_ cells (eg. *GZMA, GZMB, PRF1*). Thus, we concluded that pseudotime can correctly order cells by effectorness. Finally, we combined the pseudotime values inferred from the four trajectories into a single numeric variable corresponding to T cell effectorness (**Methods**). Interestingly, a large proportion of this variable was captured by the first UMAP component (**Figure 4G**). In conclusion, scRNA-seq enabled successful assignment of a cytokine-induced cell state and an effectorness value to each cell. Despite being separate biological variables, we hypothesized that these two axes of variation could interact to determine the transcriptional profile of each individual T cell.

### Effectorness determines the response of CD4+ T cells to cytokines

Next, we used the combined single-cell transcriptomes from all cytokine conditions (i.e. merged data set) to perform unsupervised clustering. We identified 17 clusters of T cells (**Figure 5A** and **Supplementary Table 6**) and used the trajectory analyses results to uniquely annotate each of the clusters as T cells of a given effectorness exposed to a given cytokine condition. For instance, we identified clusters of Th0, Th2, Th17 and iTreg-stimulated T_N_ cells, as well as a cluster formed of roughly equal numbers of Th17 and iTreg-stimulated T_N_ cells, characterised by high expression of TNF-signaling molecules (eg. *IL2, DUSP2, REL, TNF*) (**Figure 5A** and **Figure 5B**). Moreover, we identified four clusters of Th0-stimulated T_M_ cells, which we annotated as stimulated T_M_ cells of low, medium and high effectorness, as well as stimulated T_EMRA_. The same was true for Th17/iTreg-stimulated T_M_ cells, which localized into four groups with different effectorness (T_M low_, T_M med_, T_M high_ and T_EMRA_ cells). We also identified a group of nTreg cells, which expressed canonical markers such as *FOXP3, CTLA4* and *TNFRSF8*. This cluster contained a comparable number of cells from all the profiled cytokine conditions, suggesting that cytokines do not modify the nTreg transcriptional program. Finally, we observed a small cluster formed of T_N_ and T_M_ cells expressing high levels of IFN-induced genes, as well as a cluster characterized by high expression of heat shock proteins (HSPs) and other markers of cellular stress (**Figure 5A** and **Figure 5B**).

**Figure 5.**
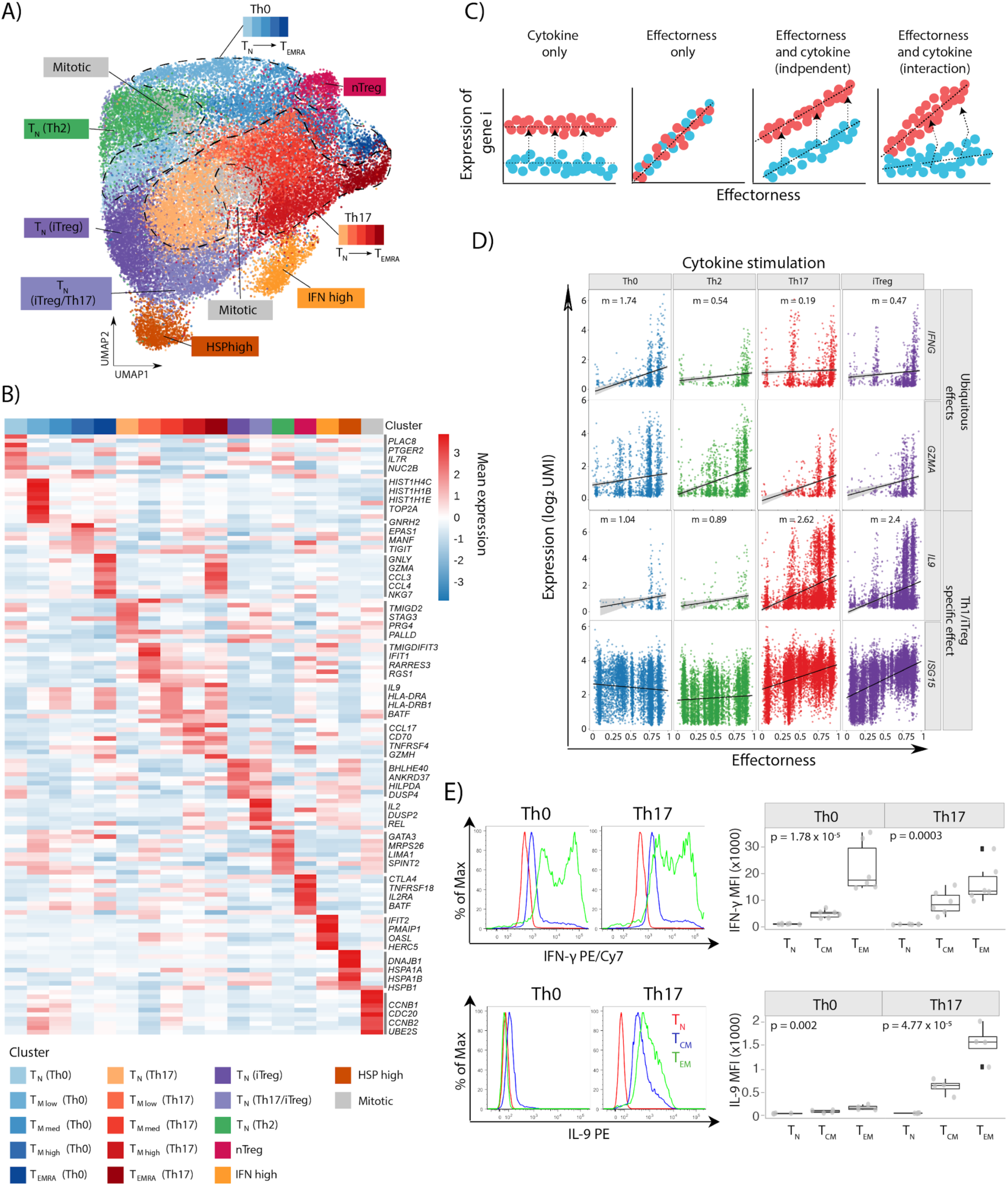
Effectorness shapes the response of CD4+ T cells to cytokines. **A)** Annotation of 17 cell clusters identified from unsupervised clustering using the top variable genes. Each cluster is annotated based on either the genes with highest expression or the effectorness and cytokine condition of the cells contained in it. **B)** Heatmap of the top 10 markers of each cluster (Wilcoxon rank sum test). Colors encode the mean expression of each gene in each cluster. Labels were added to a number of example genes for each cluster. **C)** Schematic representation of the differential expression modelled as a function of cell effectorness and cytokine conditions. Effectorness and cytokine conditions were incorporated into a linear model with interaction term (**Methods**). Genes were assigned to four groups: genes induced by cytokine-stimulation regardless of effectorness (first panel), genes which correlate with effectorness regardless of cytokine-stimulation (second panel), and genes which correlate with both effectorness and cytokine-stimulation independently (third panel), or through interaction (fourth panel). **D)** Plots of gene expression (Y axis) as a function of effectorness (X axis), with cells stratified by cytokine condition. Two example genes significantly associated with effectorness regardless of cytokine conditions (top panel) and two example genes with a strong interaction between effectorness and Th17 or iTreg-stimulation (bottom panel). Each dot represents a single cell. **E)** Levels of IFNγ and IL9 in Th0 and Th17-stimulated T_N_, T_CM_ and T_EM_ cells as assessed by flow cytometry. Representative cytometry histograms of IFNγ and IL-9 expression (upper panels) and percentages of cytokine-expressing cells in four to six biological replicates (lower panel). The p-values were calculated using one-way ANOVA.

The observed clustering of cells suggested that the nature of cytokine induced changes depends on T cell effectorness. To understand this in more detail, we modelled gene expression as a function of effectorness, cytokine stimulation and the interaction between them, where effectorness was represented as a continuous variable in the range from 0 to 1 (**Methods**). In brief, our model accounted for four possible gene expression regulatory mechanisms (**Figure 5C**): i) gene expression modulation by the presence of a cytokine irrespective of cell effectorness, ii) gene expression modulation as a function of effectorness irrespective of the cytokine condition, and gene expression modulation as a result of effectorness and cytokine-stimulation acting iii) independently or iv) jointly (interaction effect). We identified 210 genes significantly associated with effectorness (**Supplementary Table 8**). Of these, the vast majority (203 genes) were further regulated by cytokines. In particular, 12 genes showed independent effects of cytokine-stimulation and effectorness, while 191 showed an interaction effect. Within the genes with interaction effects, 12 showed an effectorness dependency only in the presence of a given cytokine, while 179 showed effectorness dependency ubiquitously (across all cytokine conditions), with the strength of this effect regulated by cytokines.

We next filtered genes by their effect sizes and identified 25 genes with a strong effectorness dependency, irrespective of cytokine stimulation. These included the costimulatory molecule *TNFRSF4* (encoding for OX40), which is known to be critical in the maintenance of memory T cell responses^41^, as well as effector molecules involved in target cell killing such as granulysin (*GNLY*), *GMZA*, *CCL3* and *IFNG* (**Figure 5D**). The expression of these genes increased proportionally to T cell effectorness. In addition, we identified 37 and 16 genes strongly associated with effectorness upon Th17 and iTreg-stimulation, respectively. These genes included cytokines like *IL2* (which decreases proportionally to effectorness upon Th17 and iTreg-stimulation) and *IL9* (which increases proportionally to effectorness for the same conditions) (**Figure 5D**). Moreover, genes induced by type I IFNs (eg. *ISG15, IFIT1, IFIT2, IFIT3*) also increased proportionally to effectorness upon iTreg and Th17-stimulation. This is in line with our observation from bulk RNA and protein expression, where we found that the type I IFN response was differentially regulated in T_N_ and T_M_ cells in response to Th17 and iTreg-stimulation (**Figure 2C**). To further validate these effects at the protein level, we isolated T_N_, T_CM_ and T_EM_ cells (**Supplementary Figure 5**). We then stimulated cells in the presence or absence of Th17 polarizing cytokines and quantified IFNγ and IL-9 expression upon restimulation (**Methods**). Our results replicated the observations from single-cell RNA-seq. Namely, the levels of IFNγ increased proportionally to effectorness in both Th0 and Th17-stimulated cells (**Figure 5E**). In contrast, the levels of IL9 only marginally correlated with effectorness in Th0 cells, but substantially increased with effectorness in Th17-stimulated cells (**Figure 5E**). This confirmed our observations from the transcriptome and suggested that key T cell functions such as cytokine secretion are under the control of both effectorness and environmental cues.

In summary, we identified a gradient of T cell effectorness which is present both before (resting state) and after stimulation. We further showed that this effectorness determines the response of single T cells to cytokines in their environment.

## Discussion

Cytokines have been extensively studied in the context of naïve T cell polarization, but the response of memory T cells to different cytokines remains understudied. Here we investigated the effects of T cell polarizing cytokines on T_M_ cells by integrating cytokine-induced changes in the transcriptome and proteome and comparing these to the responses of T_N_ cells. Our study expands our understanding of how cytokines modulate naïve and memory T cell functions. For example, we demonstrate that early changes in gene expression of both T_N_ and T_M_ cells are dominated by the response to T cell activation, while cytokine-induced changes are apparent only at the later stages of stimulation. This suggests that polarization to Th1, Th2, Th17 and iTreg occurs after the initiation of T cell activation and it fine-tunes the response of T cells. Using a combination of transcriptomic and proteomic data we defined proteogenomic gene signatures which captured well established hallmark genes induced by specific cytokines, as well as new marker genes. Both the identified genes and more broadly our dataset are valuable resources to researchers using *in vitro* cell models of cytokine polarization.

Our multi-omic data show that the response to cytokines can be strikingly different between naïve and memory T cells. While T_N_ cells respond to all cytokine polarizing conditions by acquiring a distinctive phenotype, we observed that T_M_ cells do not respond to Th2 polarization. Furthermore, we found that iTreg cells share a large proportion of their transcriptional program with Th17 cells, which is consistent with both cell states being generated in response to TGF-β^6, 8^. This is in line with previous evidence that iTreg cells can convert to Th17 in inflammatory contexts^21^. However, unlike naïve T cells, which upon iTreg-stimulation induced hallmark Treg markers such as *FOXP3* and *CTLA4*, memory T cells converge on the same cell state in response to Th17 and iTreg-stimulation. This cell state is characterized by high levels of *IL17F,* suggesting that iTreg-stimulation in memory T cells induces a Th17-like phenotype. As such, our study demonstrates that memory T cells do not acquire a regulatory phenotype upon iTreg polarization. This is particularly relevant in the context of disease, given that the number of memory T cells increases with age ^42^, potentially leading to a pro-inflammatory response to TGF-β.

In contrast, the response to IFN-β was conserved between naïve and memory T cells and, unlike Th polarization, was apparent even within 16h of stimulation. This is in line with the role of type I interferons in antiviral responses, which need to be triggered fast in order to prevent viral replication. Both naïve and memory T cells upregulated genes involved in RNAse L induction (*OAS2*, *OAS3*) that serve to degrade viral transcripts^43^, GTPase activity (*MX1*, *MX2*) to inactivate viral capsids and ribonucleoprotein assembly^44^, as well as proteins involved in cell lysis (*GZMA*, *GZMB*)^35^.

Using single cell transcriptomics, we show that CD4+ T cells form a natural progression from naïve to highly effector memory cells, which is accompanied by upregulation of chemokines and cytokines. We called this gradient, present both at the resting state and after stimulation, T cell effectorness. This suggests that, transcriptionally, memory T cell subpopulations (e.g. T_CM_, T_EM_ and T_EMRA_) are better described as stages in a continuous trajectory rather than as separate cell populations, as they have been traditionally described based on protein expression and surface markers^24^. Our results are in line with a similar trajectory, which was reported using simultaneous targeted quantification of mRNA and protein expression in single T cells^45^. Interestingly, a similar gradient is also present in innate T cells, as shown by a previous study where higher expression levels of effector molecules were negatively associated with ribosome synthesis and proliferative capacity^46^.

Importantly, the effectorness gradient described here closely recapitulates observations from immune cells isolated directly from tissues. Specifically, a previous study described the generation of memory T cells in the fetal intestine and profiled their transcriptome with single-cell resolution^47^. Cells from this study formed an equivalent trajectory, characterized by a smooth progression from the naïve to the memory state, and were accompanied by downregulation of naïve markers like *CCR7* and upregulation of cytokines like *IL32*. Thus, our results could begin to explain how memory cells in tissues adapt in response to inflammation.

Finally, we demonstrate that effectorness determines how CD4+ T cells respond to cytokines. In particular, we systematically identified genes which are regulated by cytokines in an effectorness-dependent manner, such as *IFNG* and *IL9*. Importantly, a number of these effects are also present at the protein level. Memory cells had previously been shown to upregulate IL-9 in response to TGF-β^48^, and it is known that this cytokine can also reprogram Th2 cells to an IL-9 secreting phenotype^20^. However, here we refined this observation to memory cells with high effectorness only (i.e. T_EM_ or T_EMRA_ cells). This is important given the role of TGF-β in Th17 cell biology, and especially since these cells show substantial diversity *in vivo*^23^. Our study suggests that cells with high effectorness that infiltrate tissues might start the strong responses to local cytokine environment. Future studies that use single-cell RNA-seq to profile inflamed tissues from immune disease patients will provide an opportunity to investigate these effects in greater detail directly in a disease context.

Our results demonstrate that memory cells can continue to adapt their phenotypes in response to Th17 cytokines, thus suggesting a mechanism which could generate the observed diversity. Understanding this will be key in the development of drug targets for autoimmune disease, as IL-17 and other Th17-cytokines are known to promote inflammation, for example in MS patients and animal models of disease^49–51^.

## Acknowledgements

We thank the Wellcome Sanger Institute Flow Cytometry facility, Sequencing and informatics team for their contribution to data generation and processing.

## Author contributions

GT and BS conceived and designed the project. ECG, BS and DJS carried out the experimental work. ECG and BS performed the data analysis. TIR, ES and MB performed the proteomics quantification. TIR processed and analyzed proteomics data. ECG, BS, TIR, DJS, DW, NN, JEG, CGCL, PGB, DFT, WCR, JSC and GT interpreted the results. GT, JSC and BS supervised the analysis. ECG, BS, TIR, NN, PGB, DFT, WCR and GT wrote the manuscript.

## Code and data availability

Upon acceptance we will make the raw sequencing data publicly available through the European Genome-Phenome Archive (EGA; http://ega-archive.org). The raw mass spectrometry data will be available via the Proteomics Identifications Database (PRIDE; https://www.ebi.ac.uk/pride/archive/) managed by EMBL-EBI. In the future we will host an interactive R Shiny object containing the UMAP embedding and gene expression derived from our single-cell RNA-seq data. The scripts for calculating proteogenomic signatures are available as an R package (https://github.com/eddiecg/proteogenomic).

## Methods

### Cell isolation and in vitro stimulation

Blood samples were obtained from six individuals for the bulk assays (naïve and memory T cells were isolated from three independent individuals, respectively) and from four additional individuals for the single-cell RNA-seq. All individuals were healthy males of 56.4 years of age on average (sd = 12.41 years). All human biological samples were sourced ethically and their research use was in accord with the terms of the informed consents under an IRB/EC approved protocol (15/NW/0282). Peripheral blood mononuclear cells (PBMCs) were isolated using Ficoll-Paque PLUS (GE healthcare, Buckingham, UK) density gradient centrifugation. naïve and memory CD4+ T cells were isolated from PBMCs using EasySep® naïve CD4+ T cell isolation kit and memory CD4+ T cell enrichment kit (StemCell Technologies, Meylan, France) according to the manufacturer’s instructions. T cells were then stimulated with anti-CD3/anti-CD28 human T-Activator Dynabeads® (Invitrogen) at a 1:2 ratio of beads to T cells. Cytokines were added at the same time as the stimulus (see **Supplementary Table 1** for a full list of the cytokines used with product details and exact concentrations). Cells were harvested after 16 hours and 5 days of stimulation.

### Bulk RNA-sequencing

A total of 3 × 10^5^ cells were resuspended in 500 μl of TRIzol™ and stored the material at −80°C until further processing. After samples were thawed at 37°C, 100 μl chloroform were added and samples were centrifuged for 15 min at 4°C and 10,000g. The aqueous phase was collected and mixed at a 1:1 ratio with 70% ethanol (Qiagen). RNA was isolated from this mixture using the RNeasy MinElute Kit (Qiagen), and RNA quality was assessed using a Bioanalyzer RNA 6000 Nano Chip (Agilent Technologies). All samples had an RNA integrity number (RIN) above 8.5. Finally, sequencing libraries were prepared using the Illumina TruSeq protocol and sequenced on an Illumina HiSeq 2500 platform using V4 chemistry and standard 75 bp paired-end reads.

### Proteomics

Pellets formed of up to 3 × 10^6^ cells were isolated and washed twice with PBS, dried and stored at −20 °C until protein extraction. Cell pellets were then lysed in 150 μl 0.1 M triethylammonium bicarbonate (TEAB) buffer (Sigma Aldrich) supplemented with 0.1% SDS and Halt protease and phosphatase inhibitor cocktail (100X, Thermo #78442). Pulse probe sonication (40% power, 4 °C and 20 seconds) was performed twice using EpiShear™, after which the samples were incubated for 10 minutes at 96 °C. Protein from cell lysates was quantified using the quick start Bradford protein assay (Bio-Rad) as specified by the manufacturer’s instructions. Protein samples were finally divided into aliquots of up to 100 μg. Protein aliquots were reduced with 5 mM tris-2-carboxymethyl phosphine (TCEP) buffer (Sigma Aldrich) and incubated for 1 hour at 60°C to reduce disulfide bonds. Iodoacetamide (IAA) was added to a final concentration of 10 mM and samples were incubated for 30 minutes at room temperature in the dark. Pierce Trypsin (Thermo Scientific) was then added at a mass ratio of 1:30, and samples were incubated overnight for peptide digestion. Digested protein samples were diluted to a total volume of 100 μl in 0.1 M TEAB buffer. TMT reagents (Thermo Scientific) supplemented with 41 μl anhydrous acetonitrile were added to the corresponding protein samples. After 1 hour, the reaction was quenched using 8 μl 5% hydroxylamine. Samples were then combined into a single tube and dried using a speedvac concentrator. Dry samples were stored at −20°C until fractionation. High pH Reverse Phase (RP) peptide fractionation was performed with the Waters XBridge C18 column (2.1 × 150 mm, 3.5 μm) on a Dionex™ UltiMate 3000 HPLC system. A 0.1% solution of ammonium hydroxide was used as mobile phase A, while mobile phase B was composed of acetonitrile with 0.1% ammonium hydroxide. The TMT-labelled samples were reconstituted in 100 μl mobile phase A, centrifuged and injected into the column, which operated at 0.2 ml/min. The fractions collected from the column were dried with the SpeedVac concentrator and stored at −20 °C until the MS analysis.

Liquid Chromatography-Mass Spectrometry (LC-MS) was performed using a Dionex™ UltiMate 3000 HPLC system (Thermo Scientific) coupled with the Orbitrap Fusion Tribrid Mass Spectrometer (Thermo Scientific). Dried samples were reconstituted in 40 μl 0.1% formic acid, of which 7 μl were loaded to the Acclaim PepMap 100 trapping column (100 μm × 2 cm, C18, 5 μm, 100Ӓ) at a flow rate of 10 μl/min. Multi-step gradient elution was performed at 45 °C using the Dionex™ Acclaim PepMap RSLC capillary column (75 μm × 50 cm, 2 μm, 100Ӓ). A 0.1% solution of formic acid was used as mobile phase A, and a 80% acetonitrile, 0.1% formic acid solution as mobile phase B. Precursors were selected with mass resolution of 120k, AGC 4 × 10^5^ and IT 50 ms were isolated for CID fragmentation with quadrupole isolation width of 0.7 Th. Collision energy was set at 35%. Furthermore, MS3 quantification spectra were acquired with 50k resolution via further fragmentation for the top 7 most abundant CID fragments in the Synchronous Precursor Selection (SPS) mode. Targeted precursor ions were dynamically excluded for 45 seconds.

Raw data were processed in Proteome Discoverer (v2.2) with SequestHT search engine (Thermo Scientific) using reviewed UniProt^52^ human protein entries for protein identification and quantification. The precursor mass tolerance was set at 20 ppm and the fragment ion mass tolerance was 0.02 Da. Spectra were searched for fully tryptic peptides with maximum 2 miss-cleavages. TMT6plex at N-terminus/K and Carbamidomethyl at C were used as static modifications. Dynamic modifications included oxidation of M and deamidation of N/Q. Peptide confidence was estimated with the Percolator node. Peptide FDR was set at 0.01 and validation was based on q-value and decoy database search. The reporter ion quantifier node included a TMT10plex quantification method with an integration window tolerance of 15 ppm and integration method based on the most confident centroid peak at the MS3 level. Only unique peptides for the protein groups were used for quantification. Peptides with average reporter signal-to-noise less than 3 were excluded from protein quantification.

### Single-cell RNA-sequencing

Cells were resuspended in RPMI media to obtain a single-cell suspension with high cell viability. Next, cells were stained with a live/death dye (DAPI) and dead cells were removed using fluorescence-activated cell sorting (FACS). Live cells were resuspended in PBS buffer and recounted using AOPI staining and the Nexcelom Cellometer Auto 2000 Cell Viability Counter. Finally, cells from four independent biological replicates were pooled in equal cell numbers into a single cell suspension for each condition. Cell suspensions were processed for single-cell RNA-sequencing using the 10X-Genomics 3’ v2 kit^36^, as specified by the manufacturer’s instructions. Namely, 1 × 10^4^ cells from each condition were loaded in separate inlets of a 10X-Genomics Chromium controller in order to create GEM emulsions. The targeted recovery was 3,000 cells per condition. Emulsions were used to perform reverse transcription, cDNA amplification and RNA-sequencing library preparation. Libraries were sequenced on the Illumina HiSeq 4000 platform, using 75 bp paired-end reads and loading one sample per sequencing lane.

### Flow cytometry

Cells were washed with FACS buffer (PBS buffer supplemented with 1% FCS and 1 mM EDTA) by centrifugation and stained with the respective antibodies. Reactions were incubated for 30 minutes at 4°C. Following two washes with FACS buffer, samples were resuspended in 200 μl of FACS buffer and data was acquired using a Fortessa analyser (BD Bioscience). All data were processed with FlowJo (v9.9, TreeStar).

### Intracellular cytokine staining

CD4+ T cells were obtained from PBMCs using the EasySep human CD4+ T cell enrichment kit (StemCell Technologies, Meylan, France). Next, CD4+CCR7+CD45RA+ (T_N_), CD4+CCR7+CD45RA-(T_CM_) and CD4+CCR7−CD45RA− (T_EM_) cells were isolated from CD4+ T cells via fluorescence activated cell sorting (FACS) using a MoFlo XDP cell sorter (Beckman Coulter) (**Supplementary Figure 5**) and polarized to the Th0 and Th17 phenotypes as described above. After five days of stimulation, activated naive and memory T cells were restimulated with 50 ng/ml phorbol 12-myristate 13-acetate (PMA) (Sigma) and 1 μM Ionomycin (Sigma) for five hours in the presence of 10 μg/ml of Brefeldin A (Sigma) at 37°C. After five hours, cells were fixed and permeabilized using the eBioscience™ Foxp3/Transcription Factor Staining Buffer Set (Thermo Fisher Scientific), according to the manufacturer’s instructions. Cells were resuspended in 50 μl of permeabilization solution and stained for cytokines, and flow cytometry was performed.

### RNA-seq data analysis

Sequencing reads were aligned to the reference human genome using STAR^53^ (v2.5.3) and annotated using the hg38 build of the genome (GRCh38) and Ensembl (v87). Next, the number of reads mapping to each gene was quantified using featureCounts^54^ (v1.22.2). After quantification, reads mapping to the Y chromosome and the major histocompatibility complex (HLA) region (chr6:25,000,000-47,825,000) were removed from the analysis. The final result from this process was a counts table of RNA expression in each sequenced sample.

RNA counts were imported into R (v3.5.1) where normalization for library size and regularized-logarithmic transformation of counts was performed using DESeq2^55^ (v1.19.52). We identified and removed batch effects using limma^56^ (v3.35.15). Exploratory data analysis was performed using ggplot2 (v3.0.0) and the base R functions for principal component analysis. Differential expression analysis was performed with DESeq2. More specifically, pairwise combinations were performed between any two conditions of interest, usually setting either resting or Th0-stimulated cells as controls. Differentially expressed genes were defined as any genes with absolute log-fold changes (LFC) larger than 1 at a false discovery rate (FDR) of 0.05.

### Proteomics data analysis

After quantification, protein abundances were normalised in order to allow comparisons between samples and plexes (mass spectrometry batches). Namely, protein abundance values were normalized to the total abundance of the respective sample (sample-wise normalization) and then scaled to the maximum abundance of the respective protein (protein-wise scaling). Data were then imported into R, where principal component analysis was performed using all the proteins with no missing values (proteins detected in all batches and samples) with base R functions. Finally, differential protein expression was analyzed by performing pairwise comparisons between any two conditions of interest. This was done using the moderated T test implemented in limma’s eBayes function^56^. When testing for differential protein expression, only proteins detected in at least two biological replicates per condition were kept. Multiple testing correction was performed using the Benjamini-Hochberg procedure^57^. Finally, differentially expressed proteins were defined as any proteins with an absolute log-fold change larger than 0.5 at an FDR of 0.1.

### Pathway enrichment analysis

Pathway enrichment analysis was performed using proteomics and RNA-seq data. To do so, genes detected at both the RNA and protein level were identified by matching gene names. Next, genes were ranked by differential gene or protein expression, respectively, compared to either resting or Th0-stimulated T_N_ and T_M_ cells. Finally, pathway enrichment analysis was performed independently in the RNA and protein data using the Perseus software^58^ (v1.6) and the 1D-annotation enrichment method^59^. The enrichment scores indicated whether the RNAs and proteins in a given pathway tended to be systematically up-regulated or down-regulated based on a Wilcoxon-Mann-Whitney test. A term was defined as differentially enriched if it had a Benjamini-Hochberg FDR < 0.05. Results were visualized in R using the pheatmap package (v1.0.10). For visualization, only unique pathways with an absolute enrichment score higher than 0.7, an FDR < 0.05 were kept. This was restricted to terms with biological relevance and that were included in either Reactome, KEGG or CORUM^60–62^.

### Identification of cell state signatures from RNA and protein expression

The correlation between RNA and protein expression was evaluated by estimating log-fold changes (LFCs) with respect to the control (Th0) in each cytokine condition and computing the Pearson correlation between RNA and protein LFCs. This was done both sample-wise and gene-wise. Resting T cells were excluded from this analysis.

Proteomics and RNA-seq data were used jointly to identify gene signature associated with each cytokine induced cell state. First, both data sets were matched by gene name to identify a common set of genes detected at both the RNA and protein level. Next, the f-divergence cut-off index (fCI) method^63^ was used to identify genes (RNA-protein pairs) with significant evidence of differential expression given their RNA counts and protein abundances. For any genes detected as significant by fCI, their normalized regularized-log (rlog) RNA counts^55^ and scaled protein abundances were used to calculate specificity scores in RNA and protein datasets, respectively. To do so, replicates from each condition were first averaged. Next, the specificity of each gene in each cytokine induced cell state was defined by normalizing the expression of each gene or protein to the Euclidean mean across its different cell states, as described elsewhere^29^.

RNA specificity was defined as:

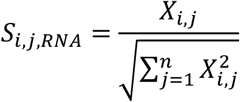

Protein specificity was defined as:

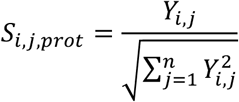

Where **X_i,j_** and **Y_i,j_** are the average RNA expression and protein abundance of gene **i** in cytokine condition **j**, respectively, and **n** is the number of cytokine conditions assessed. As the RNA expression and protein abundance are both non-negative values, S*_i,j,RNA_* and S*_i,j,prot_* are both ≥0.

Proteogenomic specificity scores were defined as the weighted sum of RNA and protein specificities for each gene:

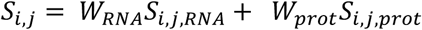

Where **S_i,j_** is the specificity score of gene i in condition j. In order to give the same weight to proteomic and transcriptomic evidence, the RNA and protein weights (**W_RNA_** and **W_prot_**) were set to 0.5.

To test which genes were more specific to one cell state than expected by chance, sample labels were randomly permuted and the specificity score was recalculated. Empirical P values were computed as the proportion of times the observed specificity score of a gene in a given cell state was larger than the corresponding permuted value. P values were corrected for the number of genes tested using the Benjamini-Hochberg procedure^57^. A total of 10,000 permutations were performed. Finally, proteogenomic signatures for each cytokine condition were defined as any genes with a specificity score larger than 0.7 and an FDR-adjusted P value lower than 0.1. This analysis was performed separately for naïve and memory T cells. The functions used to derive proteogenomic signatures are publicly available as an R package on GitHub (https://github.com/eddiecg/proteogenomic).

### Single-cell RNA-seq data analysis

Single-cell RNA-sequencing data were processed using the Cell Ranger Single-Cell Software Suite^36^ (v2.2.0, 10X-Genomics). Namely, reads were first assigned to cells and then aligned to the human genome using STAR^53^, using the hg38 build of the human genome (GRCh38). Reads were annotated using Ensembl (v87). Gene expression was then quantified using reads assigned to cells and confidently mapped to the genome.

Because each of the samples consisted of a pool of four individuals, natural genetic variation was used to identify which cells corresponded to which person. A list of common genetic variants was collected, defined as any SNP included in gnomAD^64^ with a minor allele frequency higher than 1% in the Non-Finish European (NFE) population. Next, cellSNP Cardelino v0.99)^65^ was used to generate pileups at these SNPs, resulting in one VCF file per sample. This information was then used by Cardelino^65^ (v0.99) to infer which cells belong to the same individual. Any cells which remained unassigned (with less than 0.9 posterior probability of belonging to any individual) or were flagged as doublets were discarded. In general, over 85% of cells were unambiguously assigned to an individual (**Supplementary Figure 3A**). This analysis was performed separately for each sample. To identify which individual from a given sample corresponded to an individual in a different sample, results from Cardelino were hierarchically clustered by genotypic distances between individuals. Clustering separated genotypes into four distinct groups, each group corresponding to one of the profiled individuals.

Results from RNA quantification and genotype deconvolution were imported into R and analysed using Seurat (v2.3.4)^66^. Cells with less than 500 genes detected or with more than 7.5% mitochondrial genes were removed from the data set. Counts were normalized for library size and log-transformed using Seurat’s default normalization parameters. Next, a publicly available list of cell cycle genes^67^ was used to perform cell cycle scoring and assign cells to their respective stage of the cell cycle. Cell cycle, as well as any known sources of unwanted variation (mitochondrial content, cell size as reflected by UMI content, biological replicate and library preparation batch) were regressed using Seurat’s built-in regression model. Highly variable genes were identified using Seurat and used to perform principal component analysis. The first 30 principal components were used as an input for SNN clustering and for embedding using the uniform manifold approximation and projection (UMAP)^37^. Marker genes for each cluster were identified computationally using the Wilcoxon rank sum test implemented in Seurat. Multiple testing correction was performed using FDR. Cell cycle genes were excluded from this analysis. Moreover, marker genes were required to be expressed by at least 10% of the cells in the cluster at a minimum fold change of 0.25. A total of 5 clusters were identified in resting cells and 17 clusters were found in stimulated cells. Clusters were manually annotated according to their gene expression pattern, the cytokine which cells in the cluster were exposed to and the presence or absence of hallmark genes compiled from the literature.

UniFrac distance analysis^39^ was used to test if cells exposed to two different cytokine conditions tended to form the same clusters. Pairwise UniFrac distances were computed for all combinations of cytokine conditions using all the cells captured for the respective conditions. The R package scUnifrac (v0.9.6)^40^ was used as it was specifically adapted to deal with scRNA-seq data. All parameters were set to the default values (1,000 permutations, nDim = 4, ncluster = 10).

### Pseudotime ordering and effectorness analysis

Cells were ordered into a branched pseudotime trajectory using Monocle (v2.12.0) and restricting the analysis to the highly variable genes identified by Seurat. This was done separately for each cytokine condition (resting, Th0, Th2, Th17 and iTreg), including both T_N_ and T_M_ cells. This resulted in five cytokine-specific pseudotime trajectories. Monocle was used to test for a significant correlation between gene expression and pseudotime in each trajectory. A gene was defined as significantly associated with pseudotime if its estimated q value was lower than 0.01.

The four pseudotime trajectories derived from cytokine-stimulated T cells (Th0, Th2, Th17 and iTreg) were combined into a single numeric variable. To do this, he pseudotime values of cells within each condition were scaled to the range [0,1] and combined the cells into a single data set. Finally, the association between gene expression, effectorness and cytokine-stimulation was tested with the lm() function from base R. The expression of each gene was modelled as a linear function of T cell effectorness (a numeric variable in the [0,1] range) and cytokine-stimulation (a categorical variable with levels Th0, Th2, Th17 and iTreg). An additional term was incorporated which accounted for potential interactions between these two variables, as specified in the following equation:

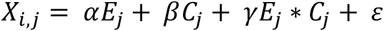

Where **X** is the expression of gene **i** in cell **j** (log2 of normalized UMIs), **E** the effectorness of cell **j**, **C** the cytokine cocktail cell **j** was exposed to and **ε** a random error term, which was assumed to follow a normal distribution with a mean of zero. The regression coefficients for effectorness, cytokine stimulation and the effectorness-cytokine interaction were represented, respectively, by **α**, **β** and **γ**. An estimate and a P value were derived for each of these coefficients in each tested gene. P values were corrected for the number of genes tested using the Benjamini-Hochberg procedure^57^. This analysis was restricted to the top variable genes identified by Seurat. All cells with zero-expression for a given gene were omitted. A coefficient was defined as significant if its corresponding FDR-adjusted P value was lower than 0.05.

## Supplementary Figures

**Supplementary Figure 1.**
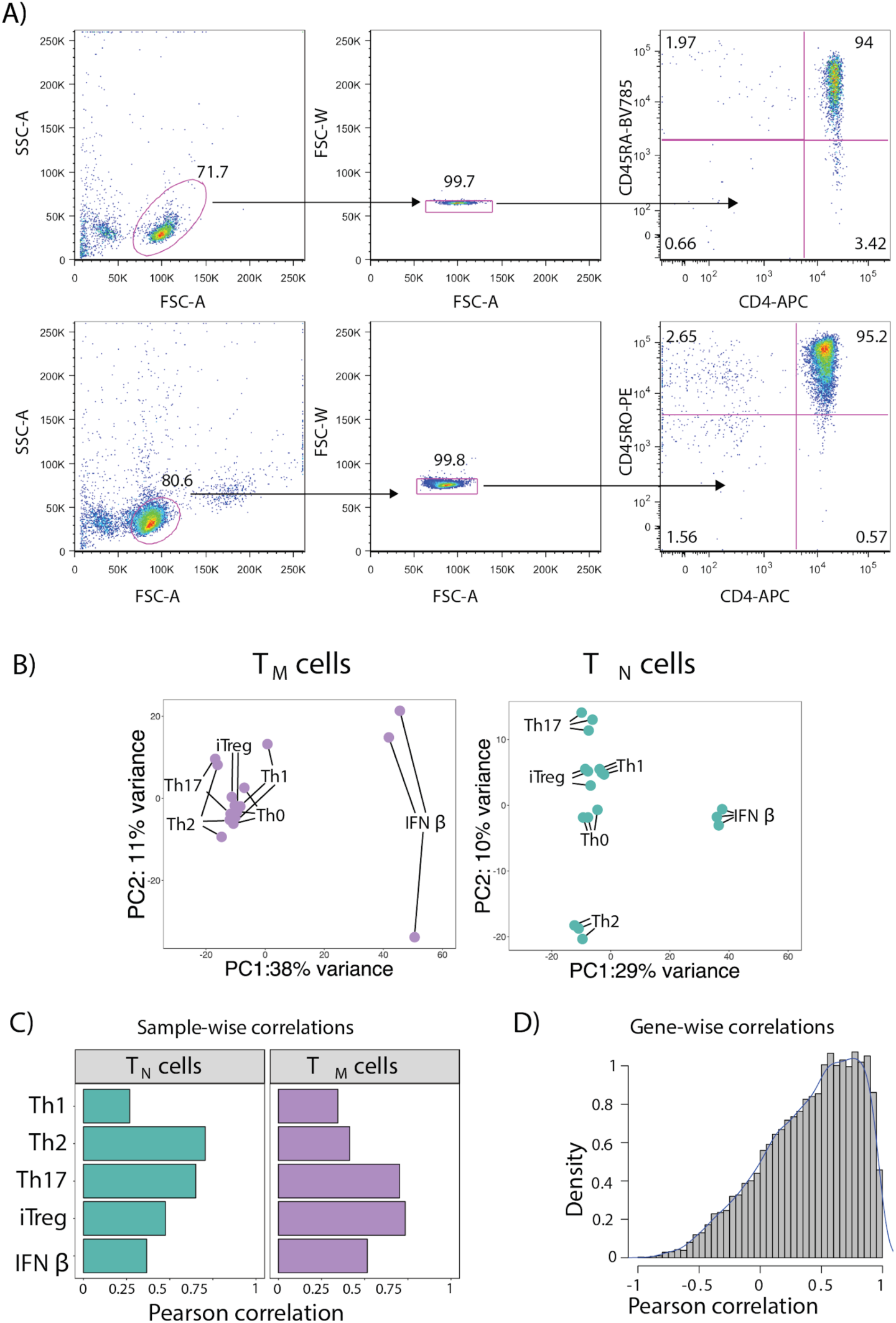
PCA and correlation between RNA and protein expression. **A)** Representative flow cytometry plots from three biologically independent samples. **B)** PCA plots from the full transcriptome and proteome of T_N_ and T_M_ cells following 16 hours of cytokine stimulations. Only stimulated cells were included in this analysis. **C)** Pearson correlations between RNA and protein log-fold changes for all genes within each cytokine condition in T_N_ and T_M_ cells. **D)** Pearson correlations between RNA and protein log-fold changes for each gene across all cytokine conditions and cell types. Resting cells were excluded from this analysis.

**Supplementary Figure 2.**
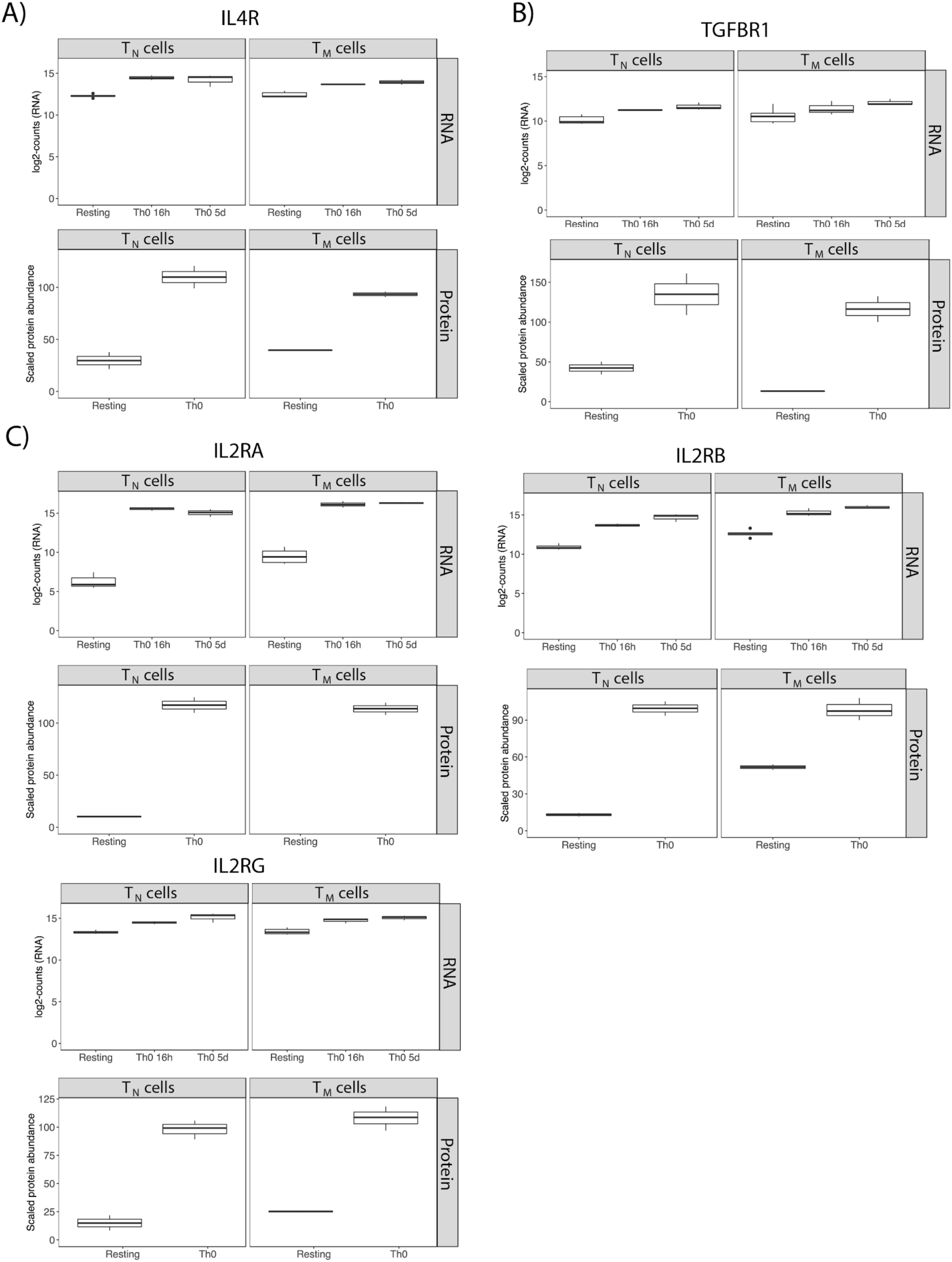
Expression of cytokine receptors. RNA levels and protein abundances of cytokine receptor components in resting and stimulated T_N_ and T_M_ cells. Subunits of the **A)** IL-4, **B)** TGF-β and **C)** IL-2 receptors.

**Supplementary Figure 3.**
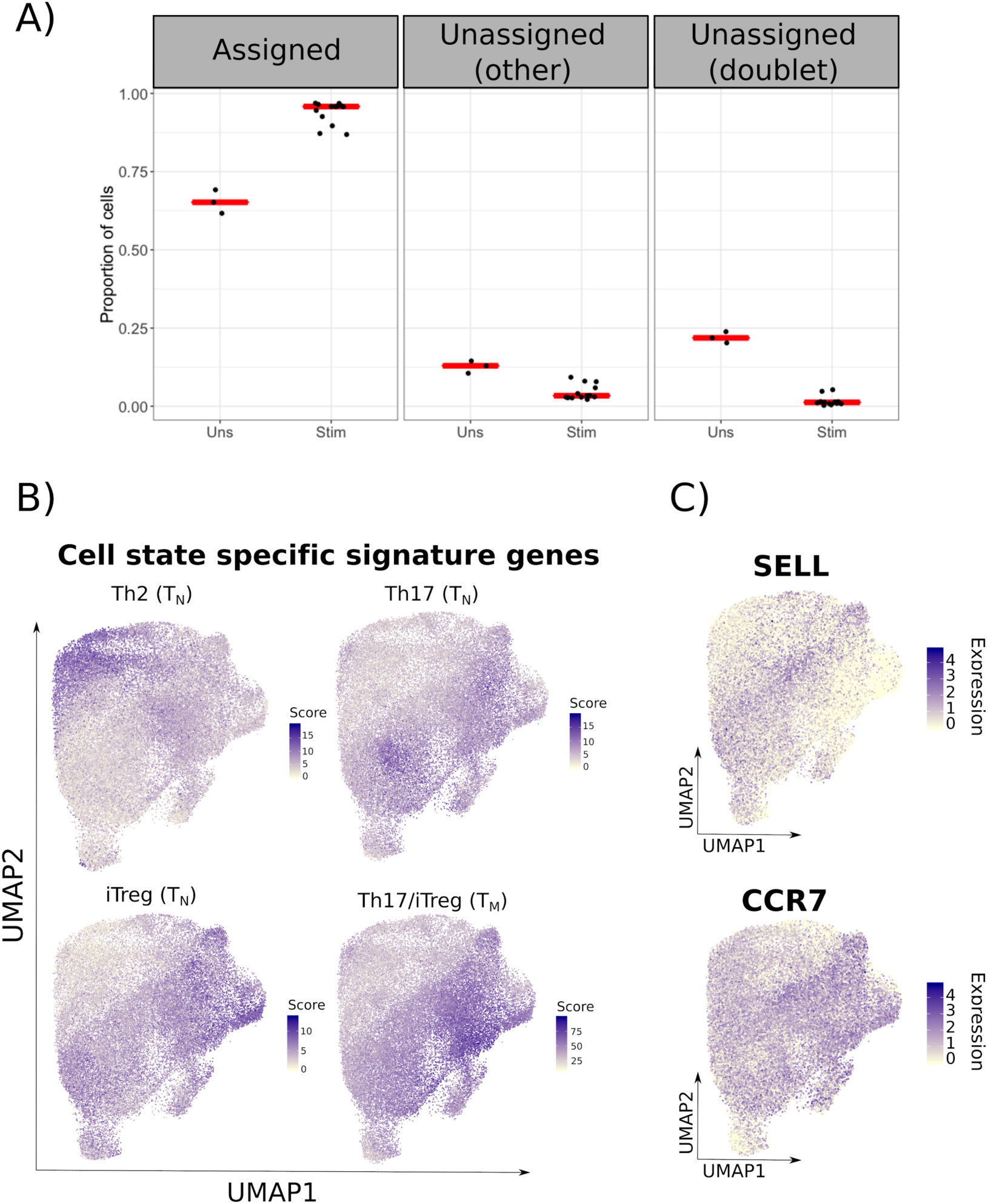
Single-cell expression of signature genes and markers in stimulated CD4+ T cells. **A)** Percentage of cells uniquely assigned to one individual (left panel), unassigned because of low posterior probability (central panel) or containing DNA from more than one genotype (doublets, right panel). **B)** UMAP embedding of stimulated T_N_ and T_M_ cells from all cytokine conditions. Colors represent the average expression of all genes in the cell state specific signatures defined from RNA and protein data **C)** UMAP embedding of stimulated T_N_ and T_M_ cells from all cytokine conditions. Colors represent the expression of *SELL* and *CCR7*.

**Supplementary Figure 4.**
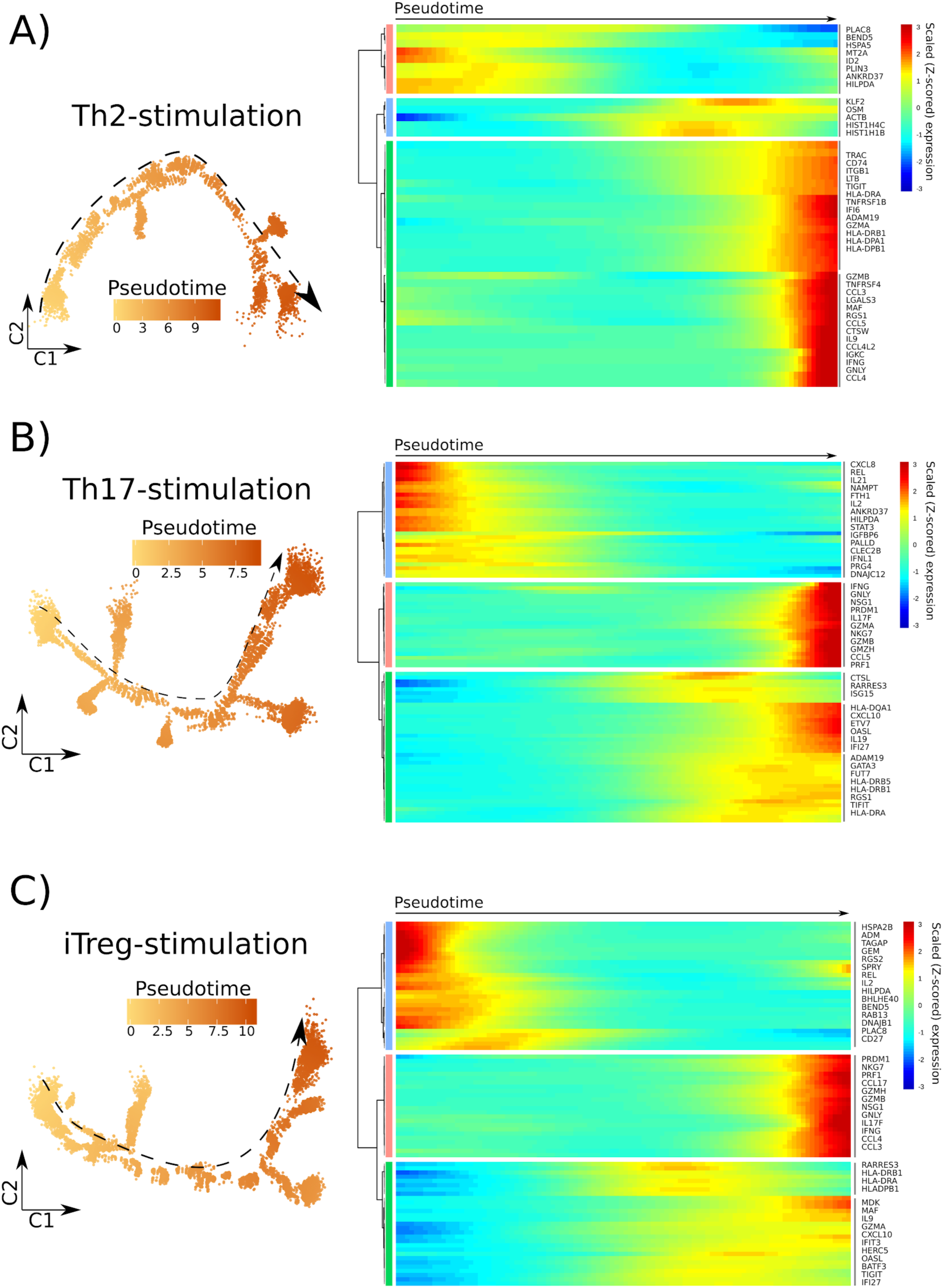
Cytokine-specific pseudotime trajectories. T_N_ and T_M_ cells ordered in branched pseudotime trajectories. Plots are colored by pseudotime value (right panels) and accompanied by a heatmap of the most variable genes along the pseudotime trajectories (right panels). Heatmap colors correspond to the scaled (Z-scored) expression of each gene in each cell. Panels correspond to four independent trajectories from **A)** Th2, **B)** Th17 and **C)** iTreg-stimulation. Labels were added to a number of example genes.

**Supplementary Figure 5.**
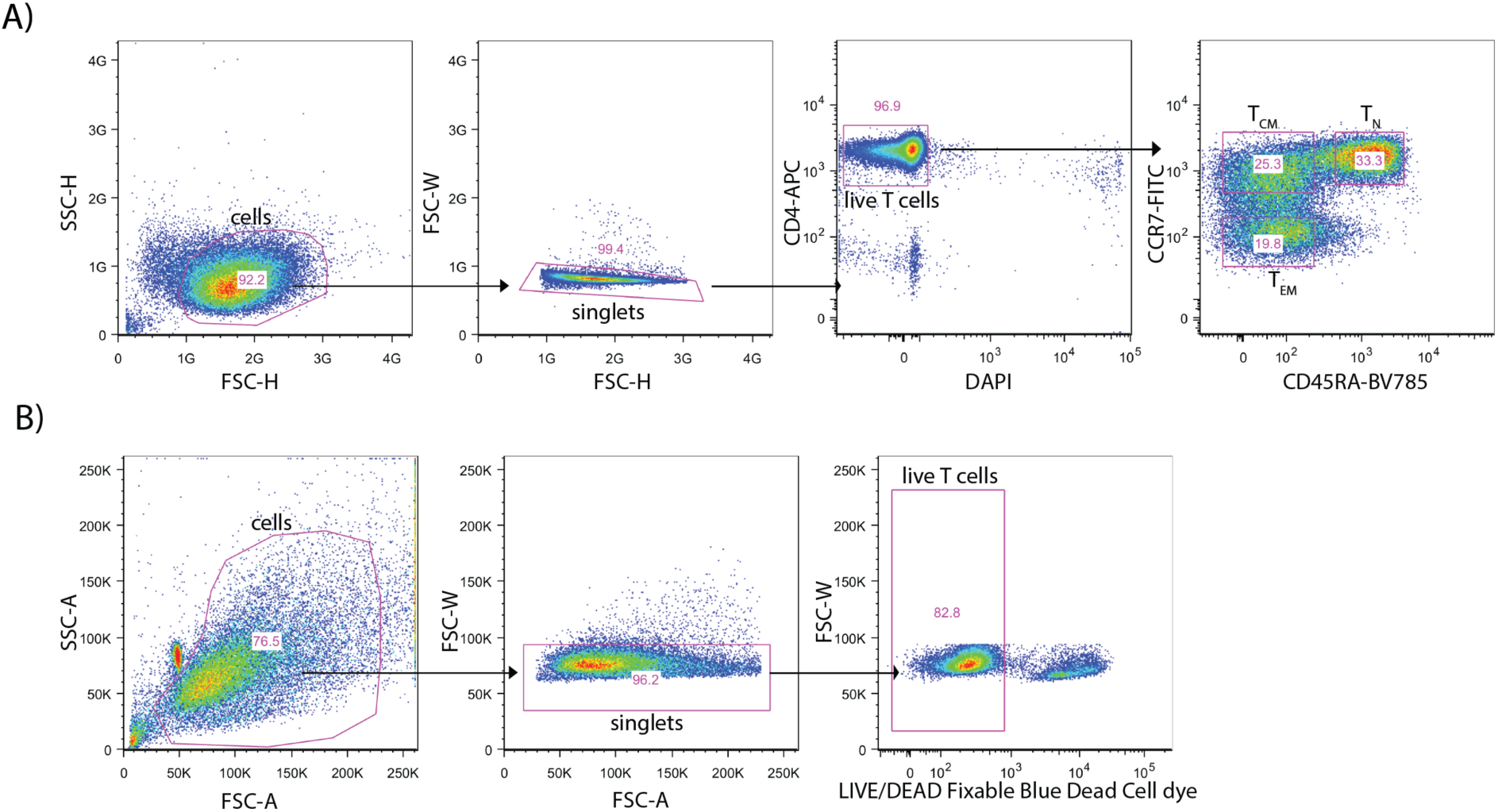
Sorting and gating strategy for cytokine staining. **A)** CD4+CCR7+CD45RA+ (T_N_), CD4+CCR7+CD45RA-(T_CM_) and CD4+CCR7−CD45RA− (T_EM_) cells were isolated from CD4+ T cells via FACS using a MoFlo XDP cell sorter. Representative flow cytometry plots of six biologically independent samples. **B)** Representative flow cytometry plots of six biologically independent samples show the gating strategy for live cell isolation after restimulation.

